# Race-dependent variability in the human tumor mycobiome

**DOI:** 10.1101/2024.06.01.596924

**Authors:** Dan Coster, Thomy Margalit, Ben Boursi, Ron Shamir

**Author notes:** equal contribution (Ben Boursi, Ron Shamir).

## Abstract

**Introduction:** Recently, Narunsky-Haziza et. al. showed that fungi species identified in a variety of cancer types may have prognostic and diagnostic signficane. We used that data in order to better understand the effects of demographic factors (age, sex, BMI, and race) on the intratumor mycobiome composition.

**Materials and Methods:** We first tested the data in view of recent critiques of microbiome data processing procedures, and concluded that the batch correction and transformation used on it may produce false signals. Instead, we explored 14 combinations of data transformation and batch correction methods on data of 224 fungal species across 13 cancer types. Propensity scores were utilized to adjust for potential confounders such as histological type and tumor stage. To minimize false outcomes, we identified as positive results only those fungi species that showed significant difference in abundance across a demographic factor within a particular cancer type, using data normalized according to all 14 combinations.

**Results and Discussion:** We observed significant differences in fungal species abundance within tumors for certain demographic characteristics. Most differences were among races in specific cancers. The findings indicate that there are intricate interactions among the mycobiome, cancer types, and patient demographics. Our study highlights the need for accounting for potential confounders in order to further understanding of the mycobiome’s role in cancer, and underscores the importance of data processing techniques.

## Introduction

The role of the mycobiome, or fungal community, in cancer development and treatment has gained appreciation recently^1^. Fungi can influence tumor biology through various mechanisms, such as the secretion of bioactive metabolites and the modulation of host immunity.^2^ Several studies have shown the role of the mycobiome in oncogenesis and its effect on immunity in pancreatic cancer.^3,4^ Other studies identified that specific fungi, such as *Malassezia*, are associated with the onset and progression of certain tumors, including colorectal carcinoma and pancreatic ductal adenocarcinoma^1,5^. Furthermore, *Aspergillus rambellii* was associated with the promotion of cancer cell growth in colorectal cancer (CRC). ^6^

Moreover, fungi show promise as a target for cancer treatment. Fungal dysbiosis has been linked to CRC, suggesting that modulating the fungal community could be an effective treatment strategy^7^. In addition, ablation of the mycobiome was reported to be protective against tumor growth in slowly progressive and invasive models of pancreatic ductal adenocarcinoma^4^. These findings highlight the potential of the mycobiome as a therapeutic target in cancer.

The impact of demographic and clinical factors on the bacterial and viral components of the tumor microbiome has been thoroughly investigated^8–10^, but little is known about their association with the cancer mycobiome. In a multicohort analysis it was shown that enteric mycobiota was altered in CRC, and that variation of fungal communities among cohorts was significantly associated with disparities in population, such as continent, sex and disease stage^6^. However, to the best of our knowledge, the effect of demographic factors on the variability of fungal species in cancer was not analyzed systematically to date. Comprehending the ramifications of these factors can improve disease understanding and treatment. The majority of studies assessing the tumor mycobiome to date have been conducted on relatively small and homogenous sample populations.^11,12^ This underscores the need for a comprehensive exploration of the impact of various factors to ensure the validity of conclusions drawn from such analyses.^11,12^

Recently, Narunsky-Haziza et al. characterized the tumor mycobiome in 17,401 tumor samples covering 35 cancer types ^13^. This work followed the extensive study of bacterial and viral microbiomes across various cancer types by Poore et al.^14^. We wished to study the demographic mycobiome variability using the data of ref^13^. However, that data was subject to some of data processing methods employed by Poore et al., which were subsequently criticized^15^. Briefly, three issues were identified: Insufficient filtering of human reads, identification as important of bacterial genera with no previous records of existence in human, and artificial signals resulting from improper data transformation and batch-correction process.

In this study, we set out to study the association between fungi abundance and patient characteristics in tumors, using the mycobiome data of Narunsky-Haziza et al. We first reevaluated the data in view of the methodological critiques. We show that one of the three issues persists, and propose a methodology to rectify it. This is achieved by applying multiple batch correction and normalization procedures. Then, we identify associations between various clinical and demographic factors and the abundance of fungi within the tumor. To mitigate potential confounding effects and enhance the robustness of our analyses, we employed both propensity scores and Inverse Probability of Treatment Weighting (IPTW) ^16,17^..

## Methods

### Data

We acquired normalized and read raw counts of intratumor fungal abundance, categorized by species, from 14,495 individual samples originating from The Cancer Genome Atlas (TCGA)^18^. Samples were labeled by tumor type, sex, age at diagnosis, and race, encompassing a total of 32 distinct tumor types. The dataset was available from Narunsky-Haziza at al. [NH22] (detailed information provided in **Supplementary Information 1**). We focused exclusively on 5,002 RNA-seq samples that were categorized as primary tumors (**Figure 1**). Additionally, Body-Mass Index (BMI) values were extracted for a subset of the samples from Hu et al.^19^.

**Figure 1:**
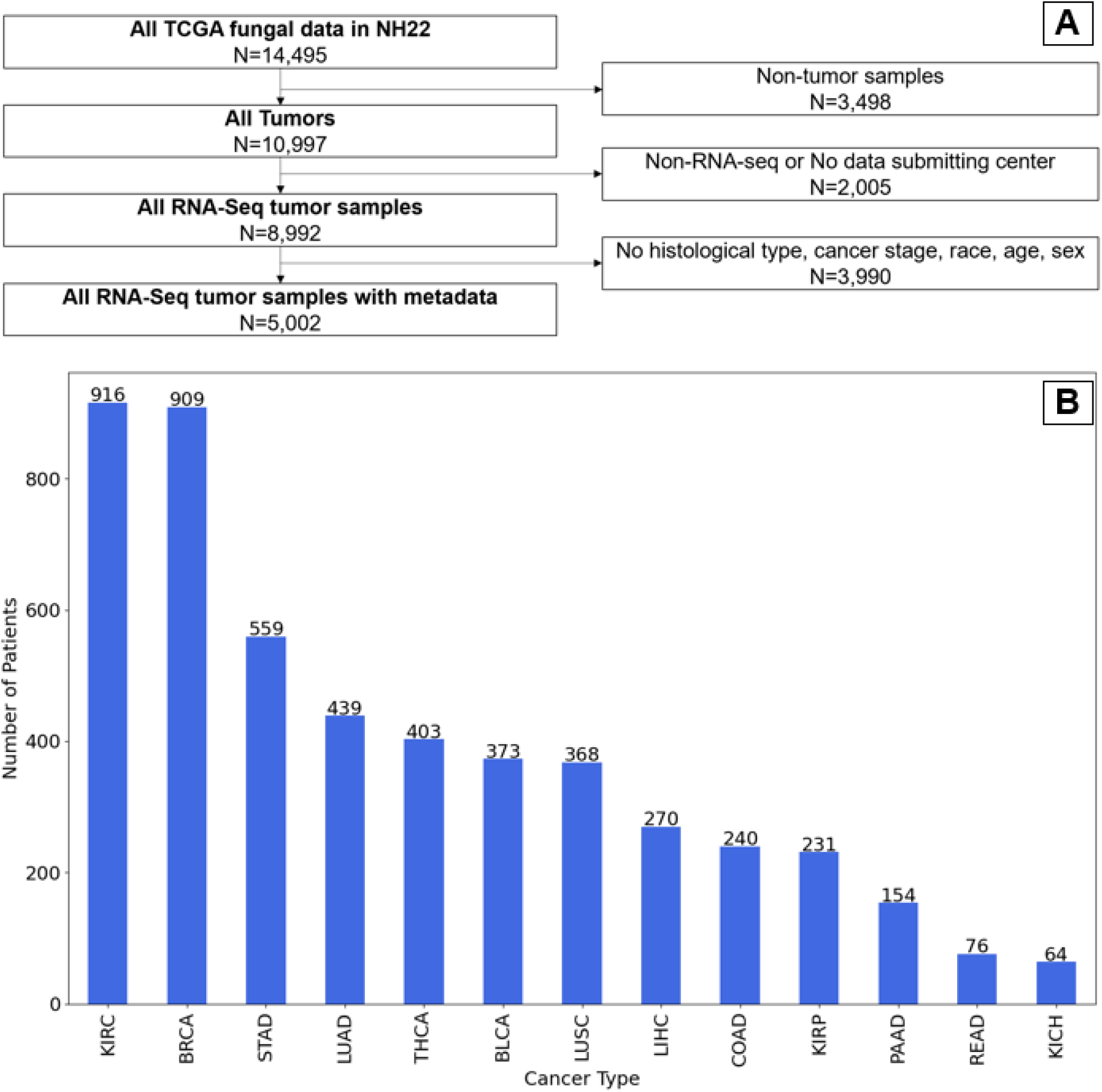
(A) Study Design. (B) The number of analyzed samples of each cancer type.

### Data Processing

NH22 employed a multistep pipeline for the retrieval of raw counts of metagenomic data. Briefly, reads deemed human or viral, too short or of poor quality were filtered out. The remaining were categorized by microbial species and underwent an additional decontamination step. Reads that mapped to a microbial reference genome were tabulated as hits. Results were summarized in a table where samples constituted the rows and microbial genome IDs the columns, containing the abundance of each species in each sample. This Decontaminated Fungal Raw Count (NH22-DFRC) table served as the basis for subsequent downstream analyses. Two additional processing steps were applied on that dataset : data transformation using *voom*^20^ (log-CPM and quantile normalization) and batch correction using *snm*^21^. This produced the Decontaminated Fungal Normalized data matrix (NH22-DFN, **Supplementary Information 1**).

### Critique of data processing techniques

NH22 used some of the same data processing methods previously employed by Poore et al. [P20], which were subsequently criticized by Gihawi et al. [G23]. Three main concerns were raised: (1) A considerable fraction of the genomic reads initially categorized as bacterial were, in fact, human. This occurred due to inaccuracies in the genomic database and the computational methodologies employed. (2) The data contained artificial signals, where bacterial genus with almost no reads in the data were deemed important for cancer classification. This happened due to the transformation methods, which were based on inappropriate statistical assumptions. (3) Some genus identified by the models as important had no previous records of existence within the human microbiome. The latter two issues were also highlighted in an earlier study^22^. In response, the authors of P20 asserted that their analytical approach was refined in their subsequent study of NH22, and that the aforementioned concerns have been adequately addressed^23,24^.

### Cohort enhancement in NH22

Narunsky-Haziza et al. expanded and improved the dataset in multiple ways: They added data from 1,183 tumor and normal tissues, which underwent additional procedures for detection of fungal contamination (WIS dataset). The TCGA data was reprocessed for better alignment and computational decontamination, and fungi species found were filtered against those found in WIS, in the Human Microbiome Projects’ gut mycobiome cohort (REF) and in a literature review of 100+ publications. Ultimately, this process led to the exclusion of 95 fungal species as potential contaminants, while retaining 224 species as bona fide non-contaminants in the dataset. In particular, this process alleviated the third concern, as only fungi previously reported in human were included in the analysis.

### Testing the number of microbial reads

We wished to test if the critique on inflated number of microbial read was true on the NH22 data. Gihawi et al. studied in detail three TCGA cancer types: bladder urothelial carcinoma (BLCA, n=683), head and neck squamous cell carcinoma (HNSC, n=334), and breast invasive carcinoma (BRCA, n=238). They reanalyzed these datasets using a stricter pipeline, thus producing novel, and lower bacterial raw counts for these cancer types (referred to as G23-BRC). To test the critique, we compared bacterial raw counts from NH22, P20 and G23, see **supplementary information 1**. To ensure an equitable comparison, only those samples and bacterial genera common to G23-BRC, NH22-BRC, and P20-BRC were analyzed, yielding data from 716 samples (BRCA, n=232, HNSC, n=330, BLCA, n=154), spanning 155 genera. Results were compared by Pearson correlation and p-values were derived by student’s t-test for correlation, and the Mann-Whitney (MW) test. Furthermore, to overcome possible sample size differences across samples, we computed the abundance ratio between pairs of genera within each sample and calculated the correlation of these pairwise ratios between samples. To avoid dependency between different ratios, we randomly subsampled sets of 77 disjoint genus pairs (covering together 154 distinct genera) and computed the correlation of their ratios. This process was repeated 2,000 times, obtaining a distribution of correlation values for each dataset.

### Testing the effect of normalization

G23 argued that the data transformation and batch-correction procedures applied in P20 (voom-SNM) generated the reported artificial signals. Since NH22 employed the same normalization methods, we wished to assess whether a similar artificial signal was present in that dataset. We focused on fungi species that were among the 10 most important features in the classifiers built in NH22 for distinguishing between tumor and normal samples in cancer types showing good classification (AUPR>0.8 and AUROC>0.9) (**Supplementary Information 1**). We sought such species with raw count of zero for more than 95% of the samples, reasoning that their contribution to the classification task should be negligible. We also compared the values of these species between the raw counts (NH22-DFRC) and the normalized dataset (NH22-DFN).

### Alternative normalizations

We used several batch-correction and transformation methods on the NH22 data instead of the voom-SNM normalization. Methods for data transformation included: Centered-Log Ratio^25^ [CLR] (with zero values offset to 1), CLR_C^26^, a variant of CLR that imputes zero counts, Relative abundance, and *voom*^20^. voom was utilized after log-CPM transformation and quantile normalization as in NH22 and P20. For batch correction we tested ComBat^27^, Batch Mean Centering^28^ [BMC], MMUPHin^29^ and PLSDA^30^. RBE^31^ produced equivalent results to BMC in our tests, so we did not include it. To avoid any potential data leakage, we selected only unsupervised batch correction methods. In our analysis, we tested all combinations of transformation and batch correction methods listed above. It is noteworthy that MMUPHin specifically accepts raw counts or relative abundance counts as input and produces the same types of data as output. Therefore, we first applied MMUPHin to our data and then the data transformation. In all other cases the transformation preceded batch correction. Additionally, since PLSDA can be used only after CLR transformation, we used it only with CLR and CLR_C. In total, we applied 14 combinations.

### Fungal differential abundance

We gathered tumor and patient characteristics from the TCGA dataset. Tumor factors included tumor stage and histological type, and patient attributes included age at diagnosis, sex, and race. Five independent variables were evaluated: Sex, age, (i.e., old (age≥70) vs. young (age < 70)), and pairwise race comparisons: European vs. Asian, European vs. African, and Asian vs. African. Our dependent variables were the fungal species.

To mitigate potential confounding effects, we employed a two-step approach. First, we computed propensity scores using logistic regression based on the independent variables. Subsequently, we applied the Inverse Probability of Treatment Weighting (IPTW) scheme as performed in ^16,17^. Categorical variables were subjected to one-hot encoding, and tumor stage was categorized into four levels (1, 2, 3, 4) to facilitate the analysis (for example 3A, 3B and 3C were considered as 3). This procedure was instrumental in correcting confounding bias. We ensured that all the potential confounders were well-balanced between the groups, with standardized differences of less than 10%, in line with previous research^17,32^. For each transformed and batch-corrected fungal specie, we fitted a linear regression model with the IPTW weights and computed the model’s p-value. Correction for multiple hypotheses was done by Bonferroni’s method. Species with P-value<0.05 were deemed statistically significant. Notably, when evaluating the effect of a patient characteristic on fungal species, we excluded it from the IPTW calculation. Thus, we repeated the IPTW procedure for each independent variable and for each cancer type with 20 or more samples in each of two groups.

Body mass index (BMI) was available for a subset of the samples. We examined its influence (binarized into obese [BMI≥30] vs. non-obese [BMI<30] categories), considering all other patient characteristics (age at diagnosis, sex, race) as potential confounders in IPTW. All data processing and statistical analyses were carried out using R statistical software version 4.2.0 and the Python programming language 3.4.

## Results

We obtained data of 14,495 samples from the TCGA and following the application of exclusion criteria (Methods) were left with 5,002 samples of tumors that had RNA-Seq data, information on the sequencing center and patient characteristics (**Figure 1A**). The number of samples per cancer type varied between 916 in Kidney Renal Clear Cell Carcinoma (KIRC) and 64 in Kidney Chromophobe (KICH) (**Figure 1B**). The distribution of demographic characteristics varied across cancer types. For example, 99% of BRCA samples were of females vs. 54% in Lung Adenocarcinoma (LUAD). In all cancer types the most prevalent race was Europeans. The distribution of the age, BMI, tumor stage and histological type also varied between cancer types (**Supplementary Table 1**).

**Table 1:** The number of fungal species that were identified as statistically significant across all 14 combinations of normalization and batch correction per each comparison.

| **<u>Cancer Type</u>** | **<u>Asian vs. European</u>** | **<u>Black vs. European</u>** | **<u>Black vs. European</u>** | **<u>Male vs. Female</u>** | **<u>Age > 70</u>** | **<u>BMI > 30</u>** |
| --- | --- | --- | --- | --- | --- | --- |
| **COAD** | - | 5 | - | 0 | 0 | - |
| **LIHC** | 1 | - | - | 0 | 1 | 1 |
| **LUAD** | - | 2 | - | 0 | 0 | - |
| **LUSC** | - | 2 | - | 0 | 0 | - |
| **STAD** | 9 | - | - | 1 | 0 | - |
| **THCA** | 0 | 0 | 1 | 0 | 0 | - |
| **PAAD** | - | - | - | 0 | 0 | - |
| **BRCA** | 0 | 0 | 0 | - | 1 | - |
| **KIRP** | - | 0 | - | 0 | 0 | 0 |
| **BLCA** | 0 | 0 | 0 | 0 | 0 | 0 |
| **READ** | - | - | - | - | 0 | - |
| **KIRC** | - | 0 | - | 0 | - | - |
| **KICH** | - | - | - | 0 | - | - |

Recently, Gihawy et al. (G23) raised concerns regarding the validity of the analysis of microbiome data in the TCGA by Poore et. al (*Nature*, 2020) (P20). As NH22 used some of the same preprocessing methods, we first wished to ensure the validity of the NH22 data, by checking these concerns on it (see Methods for further details). One concern was an inflated microbial read counts. To check this, we compared the data of NH22, P20 and the subset of data of three cancer types that was re-analyzed in G23. To ensure an equitable comparison, only samples and bacterial genera common to G23, NH22, and P20 were analyzed, yielding 716 samples (BRCA, n=232, HNSC, n=330, BLCA, n=154) spanning 155 genera. The total number of bacterial reads in NH22 was 2-3 magnitudes of order lower than in P20, and 1-2 magnitude of order higher than in G23 (**Figure 2A**). Similar ratios were noted in the distributions of the bacterial read counts per sample (**Figure 2B**). Thus, the differences between NH22 and G23 are indeed less significant than between P20 and G23. Furthermore, the read count per sample of G23 showed stronger correlation with NH22 than with P20 (*r* = 0.54 vs. *r* = 0.02, **Figure 3A, 3B**). Similar trends were observed when each cancer type was evaluated separately **(Figure 3C, 3D**).

**Figure 2:**
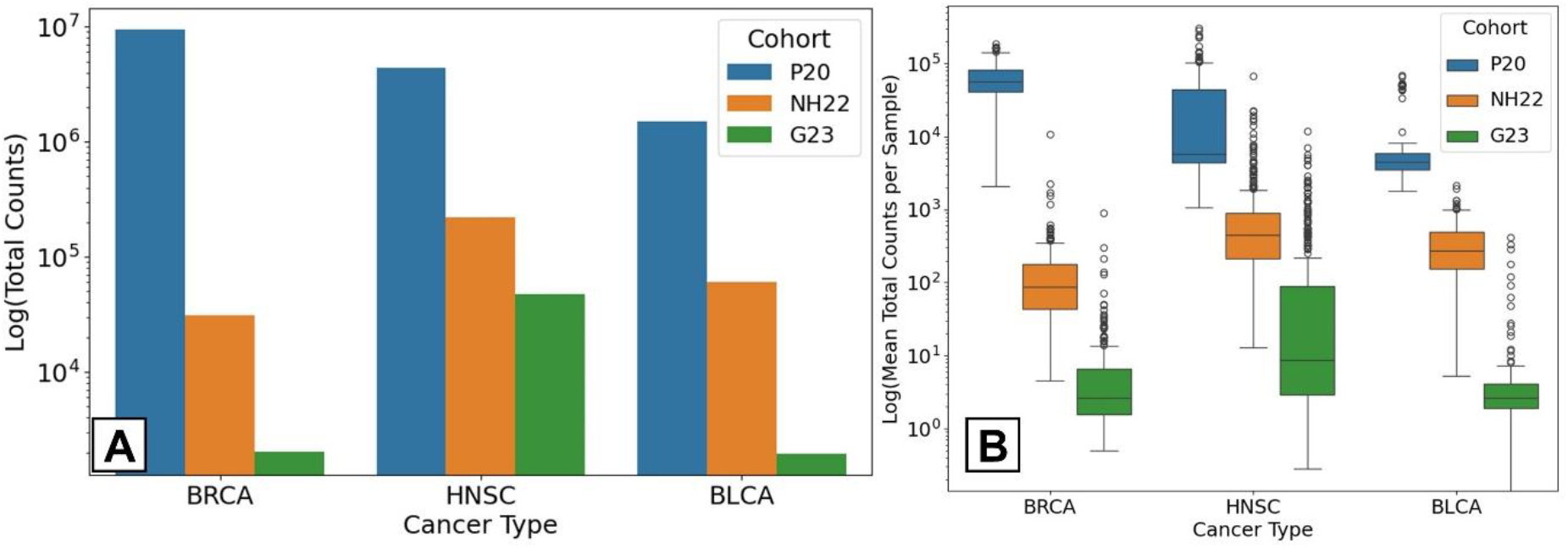
Statistics on the count of bacterial reads after filtering in the P20, NH22 and G23 datasets. (A) Total count. (B) Boxplots depicting the distribution of the counts per sample.

**Figure 3:**
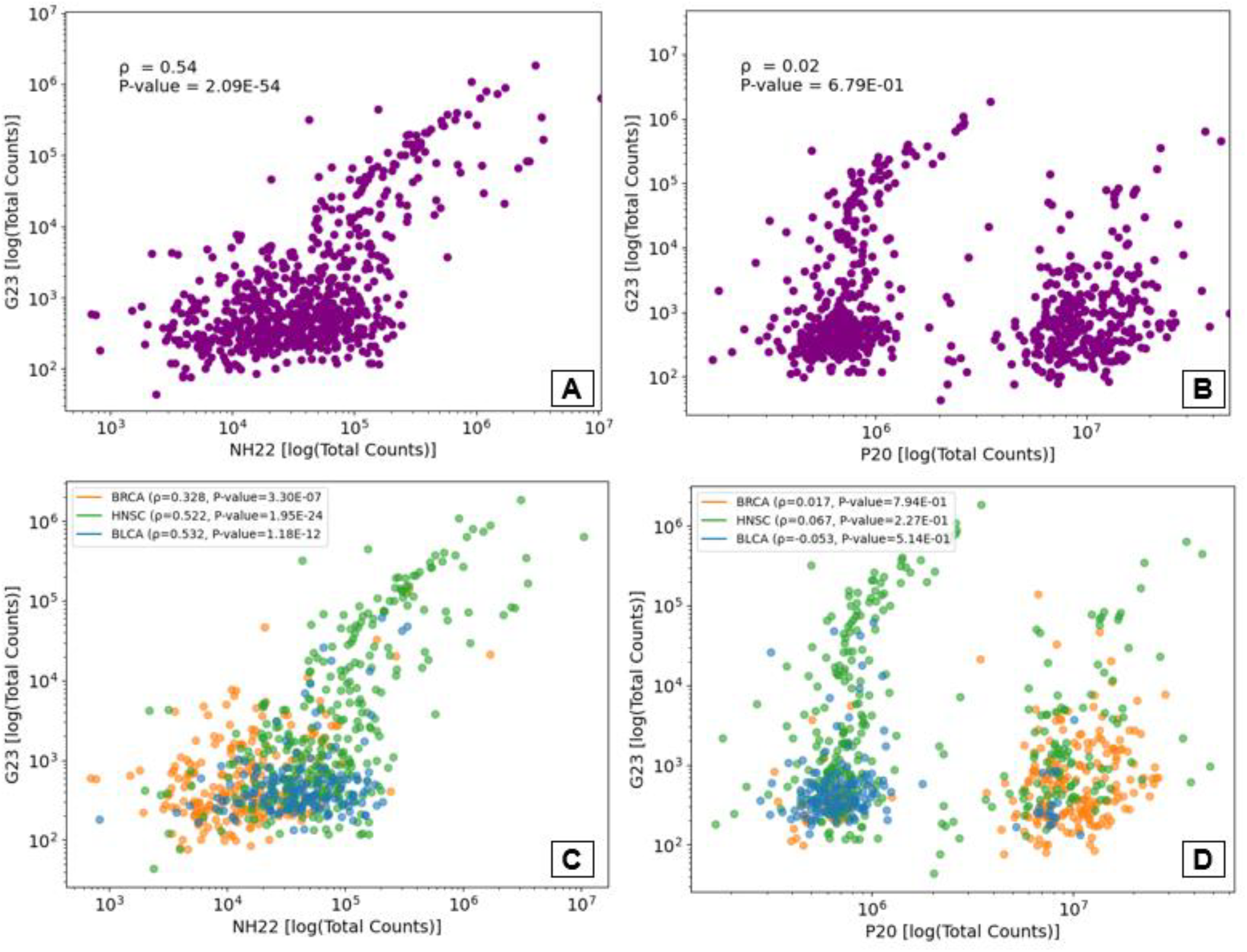
Correlation plots of the total bacterial read count per sample. (A) G23 vs. NH22. (B) G23 vs P20. (C) G23 vs. NH22, analyzed separately for each cancer type. (D) G23 vs P20, analyzed separately for each cancer type. Each point corresponds to a sample.

Next, we compared the bacterial data of NH22 and G23 at the genus level. The total number of reads per genus were correlated (*r* = 0.67, p=4.17E-21, **Figure 4A**), and this correlation persisted in two of three cancer types when analyzed separately (BRCA: *r* = 0.323, p=4.01E-05, HNSC: *r* = 0.757, p=2.83E-30, BLCA: *r* = 0.025, p=0.075, **Figure 4C**). P20 was far less correlated to G23 in this analysis (**Figure 4B, 4D**).

**Figure 4:**
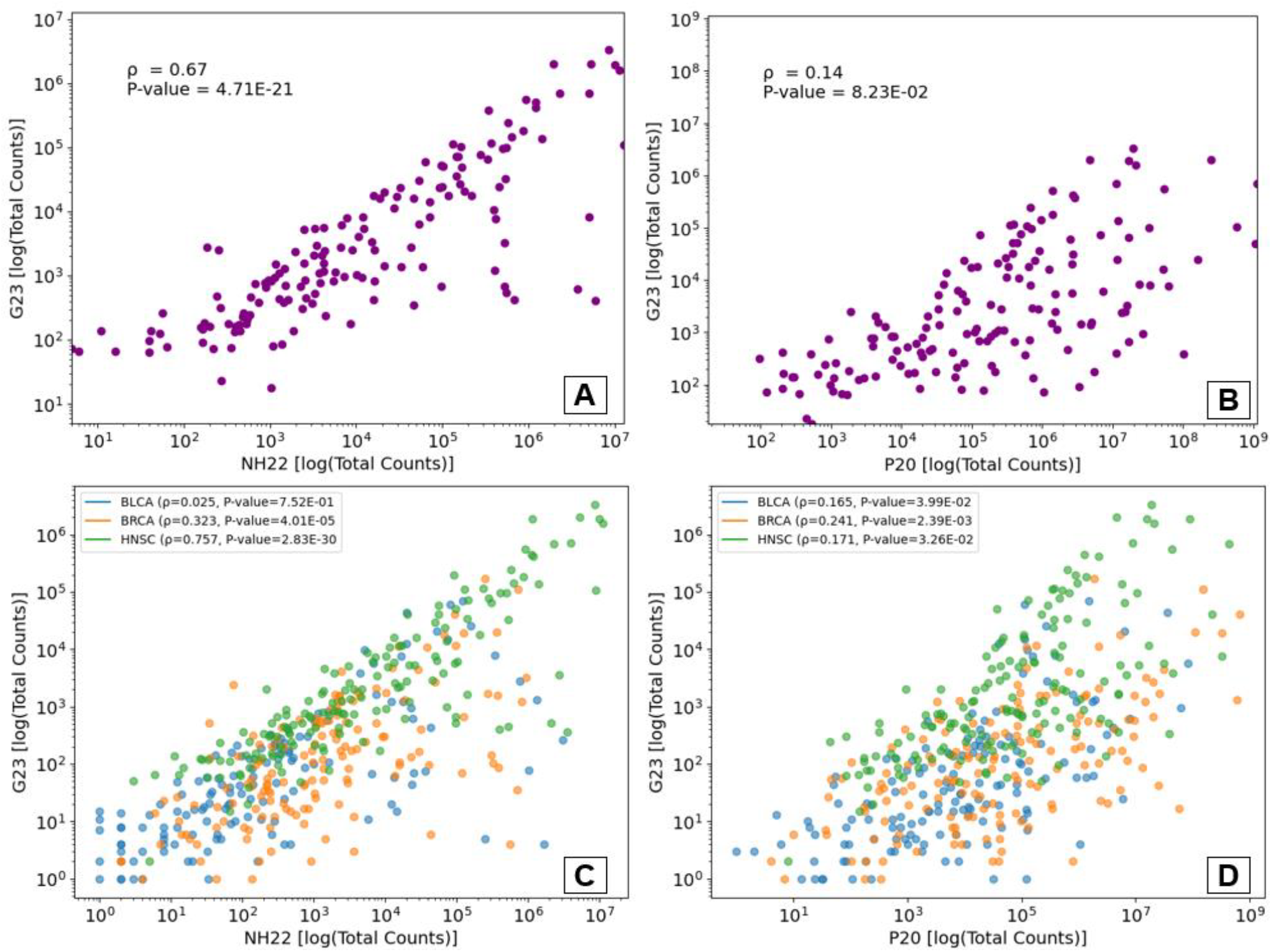
Correlation plots of bacterial read count per genus. (A) Total count, G23 vs. NH22 (B) Total count, G23 vs. P20. (C) Correlation analyzed by cancer type, G23 vs. NH22. (D) Correlation analyzed by cancer type, G23 vs. P20. Each point corresponds to a genus.

As an additional test that avoids potential differences in the total read counts among the datasets, we computed the ratios between read counts of different genera within each dataset (pairwise ratios) and then calculated the correlation between these ratios in NH22 and G23, and in P20 and G23 (taking steps to avoid dependency, see Methods). The correlation of NH22 to G23 was significantly higher than that of P20 to P23 (p=5.59E-132, **Supplementary Figure 1)**. Similar results were observed per cancer type (results not shown).

Next, for each bacterial genus and cohort, we computed the correlation between the genus’ read counts across samples, and plotted the distribution of the correlation values across the genera. The correlations between NH22 and G23 were significantly higher than between P20 and G23 (**Figure 5, Supplementary Table 2**). Analyzing separately each of the five genera with the highest number of samples with no zero read counts in G23 were (**Supplementary Figure 2**) *Pseudomonas* (NH22 vs. G23; *r* = 0.684, p=1.2E-99, P20 vs. G23; *r* = 0.02, p=0.6), *Staphylococcus* (NH22 vs. G23; *r* = 0.565, p=2.7E-50, P20 vs. G23; *r* = ™0.155, p=3.3E-05), *Acinetobacter* (NH2 vs. G23; *r* = 0.197, p=1.1E-06, P20 vs. G23; *r* = ™0.02, p=0.6), ), *Corynebacterium* (NH22 vs. G23; *r* = 0.959, p=8.8E-306, P20 vs. G23; *r* = 0.957, p=3.3E-313) and *Klebsiella* (NH22 vs. G23; *r* = 0.944, p=1.4E-171, P20 vs. G23; *r* = 0.017, p=0.7).

**Figure 5:**
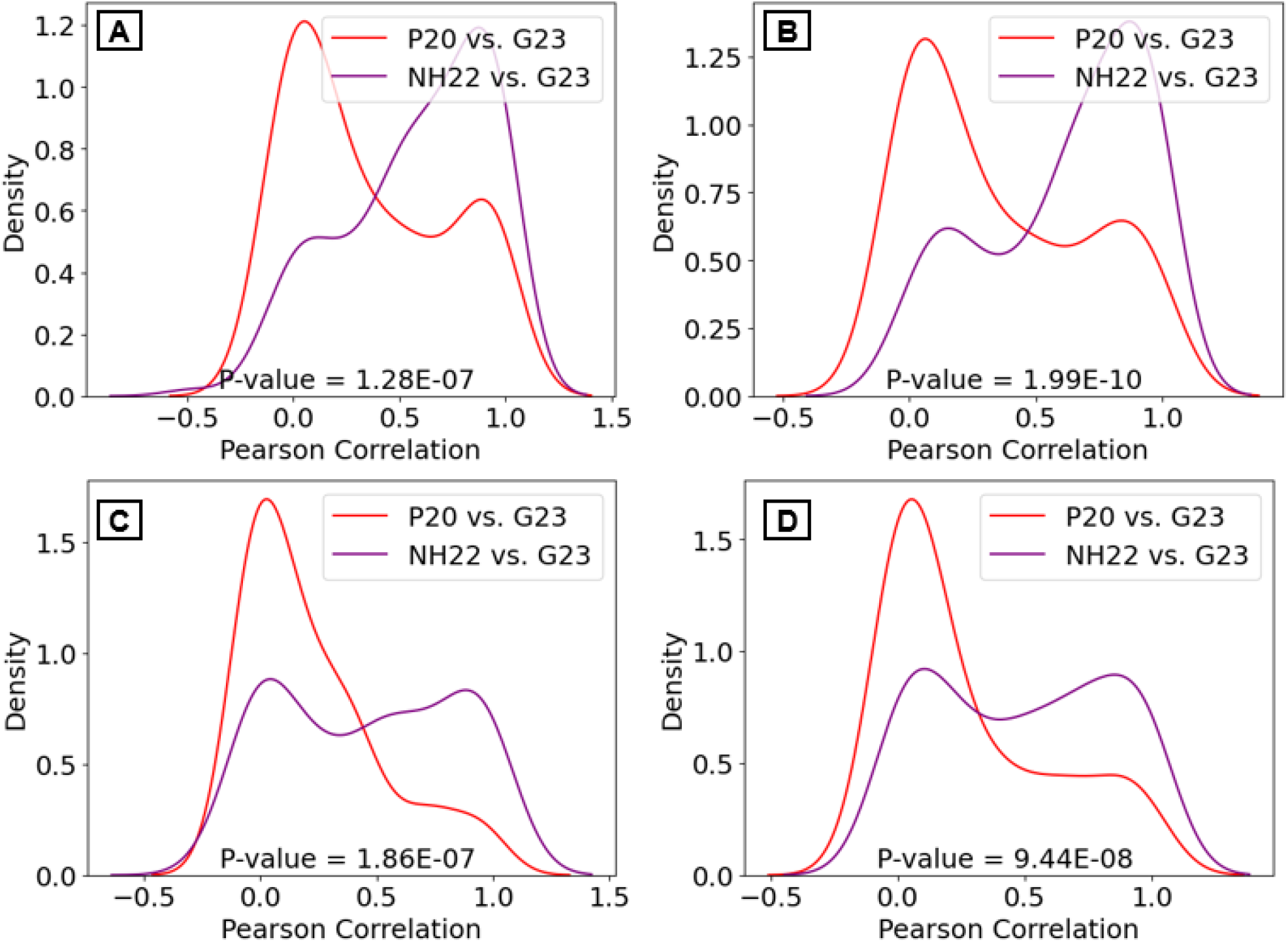
Distribution of the correlation between genera read counts across samples. The distribution includes the values for all genera. Each figure shows the correlations for NH22 vs. G23 and P20 vs. G23. (A) all cancers. (B) HNSC. (C) BLCA. (D) BRCA.

Similar results were obtained when repeating this analysis per cancer type (**Supplementary Figure 3**,**4**,**5** and **Supplementary Table 2**). Overall, the analysis results showed that the issue of inflated bacterial read count in P20 raised by G23 was addressed to a large extent in NH22, and that many characteristics of the NH22 data were similar to that in G23.

Another important issue raised by G23 in P20 was the procedure used to remove batch effects and convert the raw counts to normalized values. P20 used for that purpose Voom-SNM^20,21^. Cancer type classifiers built in P20 using the processed data included as important features bacterial genera that were in fact absent in most or all samples. Since Voom-SNM was also used in NH22, we wished to test if fungi-based classifiers built by NH22 also suffered from the same problem. We examined the ten genera with the highest feature importance used in the classifier of primary tumor vs. normal tissue, and found some examples of this phenomenon:

*Parastagonospora* was the 3^rd^ most important feature for the classifier in prostate adenocarcinoma (PRAD). Out of 666 PRAD samples of primary tumor and normal tissue (of both RNA-seq and WGS), 665 had zero *Parastagonospora* reads and only one had one read. As illustrated in **Figure 6**, the extremely non-random and right skewed distribution of the primary tumor normalized values–all of which except of one started as raw values of zero– makes it easy for a machine learning classifier to separate the PRAD primary tumor samples from normal solid tissue samples. Based on that distribution a model that splits the samples using the threshold 12.457 labels 332/351 of the positive samples correctly (PPV=94.6%), with relatively high sensitivity of 55.33%. *Ramularia* was the 6^th^ most important feature for the classifier in KIRP. Out of 345 KIRP samples, 344 had zero *Ramularia* reads and one had one read. The distributions of the normalized values are extremely different (**Supplementary Figure 6**). A threshold of 17.36 obtains PPV=95.6% with sensitivity of 49.84%. We conclude that following Voom-SNM normalization, false signals persist in the NH22 processed data.

**Figure 6:**
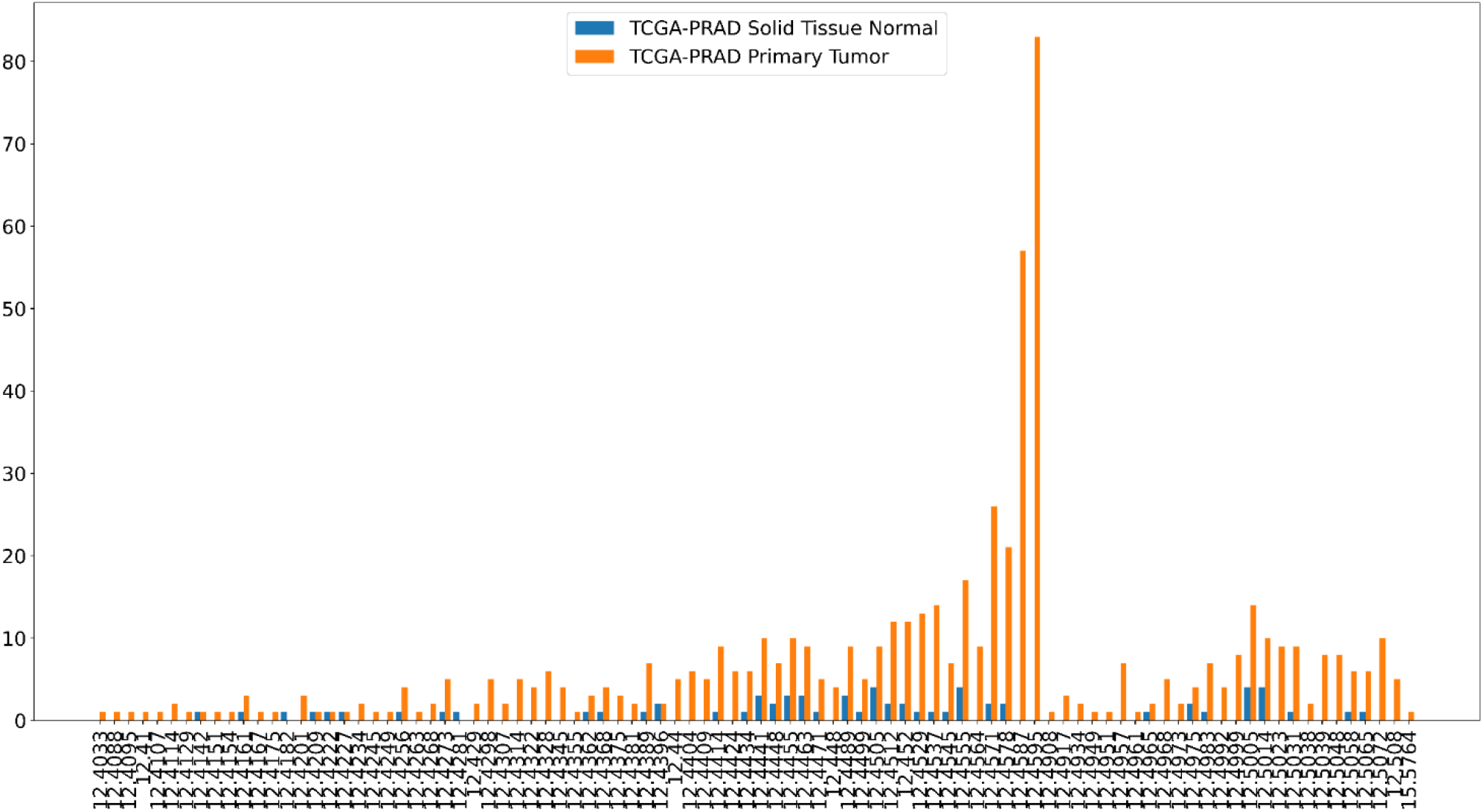
The effect of Voom-SNM normalization. Distribution of normalized counts of *Parastagonospora* for Prostate Adenocarcinoma primary tumor samples (PRAD, blue) vs. normal tissue samples (orange). Before the normalization, out of 666 samples, 665 had zero count and one had a count of 1.

Our main question in this study was whether fungi species abundance varies in cancer patients with different demographic characteristics. Batch correction and data normalization are key for the answer, so instead of Voom-SNM, we used 14 combinations of normalization and batch correction approaches (**Figure 7** and Methods). NH22 used SNM to correct two types of batch errors: specimen type (RNA-seq or DNA) and the sequencing center of origin. Since most current batch correction methods were designed to correct one type of error, we decided to use only the RNA-Seq samples, since they cover most of the cohort (8,992/10,997 tumor samples). We used propensity score and IPTW for correction of potential confounders: age, sex, race, tumor stage, and histological type (Methods). IPTW was applied to each demographic factor separately. Then, for each fungal species, a linear regression was fitted to the IPTW weights, and p-values were Bonferroni corrected. To err on the side of caution, we sought species that were significant in all 14 combinations. Twenty-four species showed consistent statistically significant differences between race groups (**Table 1**). In the comparison between European vs. African there were five significant fungal species in colon adenocarcinoma (COAD): *Candida orthopsilosis, Penicillium expansum, Diutina rugosa, Aspergillus alliaceus, Aspergillus pseudonomius*, two species in LUAD: *Diutina rugosa, Aspergillus pseudotamarii*, and two species in Lung Squamous Cell Carcinoma (LUSC): *Candida orthopsilosis, Aspergillus welwitschiae*. In the comparison between European vs. Asian there were nine significant fungal species in Stomach Adenocarcinoma (STAD): *Aspergillus clavatus, Aspergillus terreus, Aspergillus aculeatus, Penicillium digitatum, Pestalotiopsis fici, Penicillium expansum, Aspergillus novofumigatus, Aspergillus campestris* ,*Ramularia collo-cygni* and one species, *Metarhizium robertsii*, in Liver Hepatocellular Carcinoma (LIHC). In the comparison of Asian vs. African populations, one species, *Candida auris*, was found in the Thyroid Cancer (THCA). In comparison of Males vs. Females, one species, *Purpureocillium lilacinum*, was found in STAD. In comparison of BMI≥30 vs. BMI<30, one specie, *Diutina rugose*, was found in LIHC. In comparison of old (Age >70) vs. young (≤70) one species, *Hyphopichia burtonii*, was found in LIHC and one species, *Aspergillus oryzae*, was found in BRCA. As an additional test, we also compared the raw count distributions of these significant fungi. Remarkably, 21 out of 22 species demonstrated statistically significant associations for the corresponding cancer type and characteristic, even without undergoing normalization, batch correction, or adjustment for confounders (**Supplementary Table 3**).

**Figure 7:**
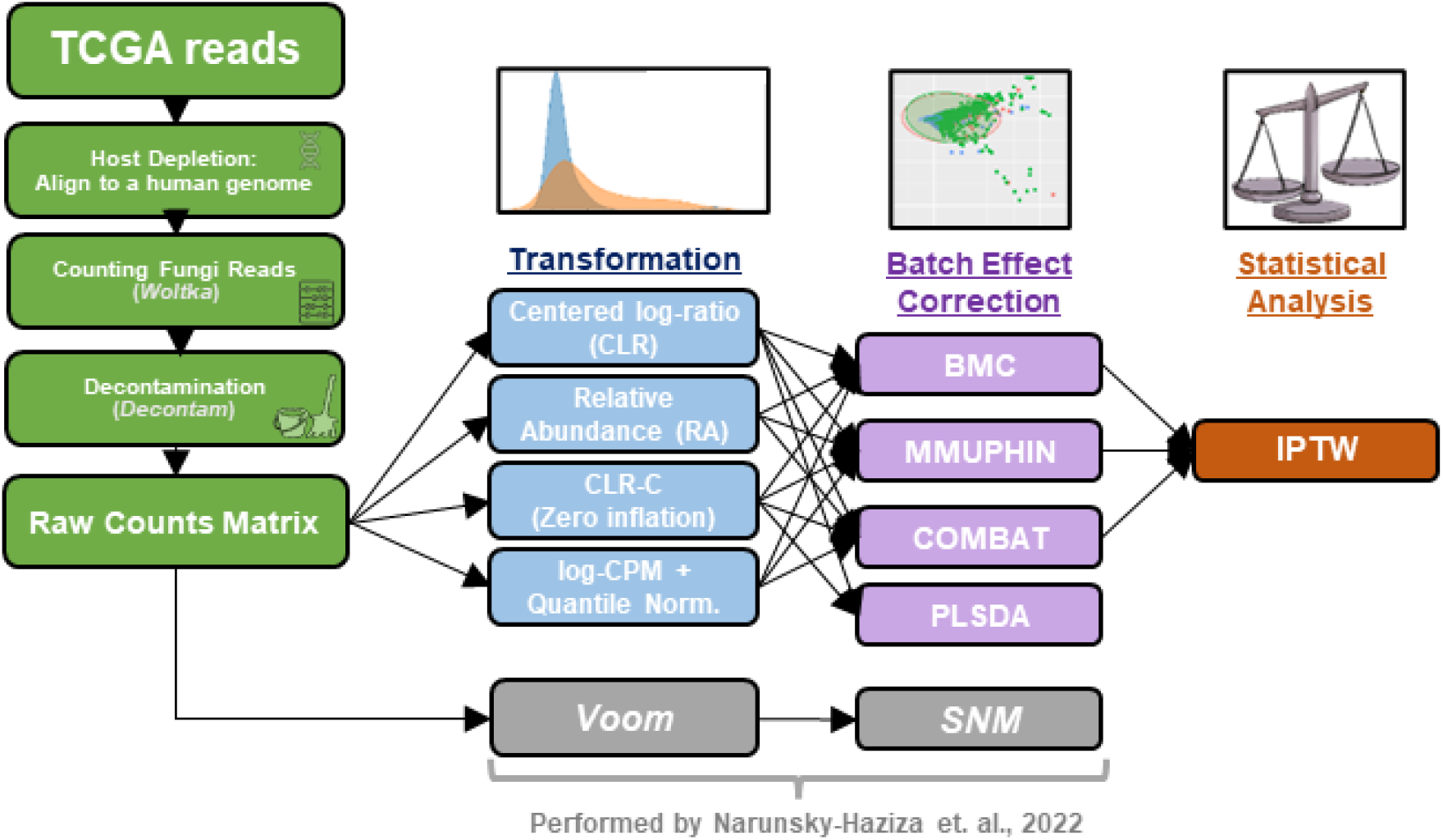
The pipeline used to clean and normalize the fungi data. The central part shows the combinations of transformation and batch correction used.

## Discussion

We aimed to investigate the effect of different demographic factors on the abundance of different fungal species within cancer tumors. By analyzing a dataset comprising over 5,000 tumor samples from the TCGA, we revealed distinct patterns of fungal abundance associated with race, gender, age, and obesity, highlighting the complex interplay between demographic factors and the tumor microenvironment. Differences across races were mostly observed. These findings underscore the importance of considering demographic diversity in cancer research, particularly in the context of the intratumor mycobiome. By accounting for such factors, can help researchers to understand the underlying cancer mechanisms better.

We took several steps to check recent concerns raised about microbial analyses in cancer. In particular, we developed a pipeline that utilizes a consensus of multiple methods for batch correction and data normalization and also utilized propensity scores when calling association between a particular fungus and a particular demographic factor in a certain cancer. This strategy increases robustness and reliability of the results and reduces the risk of spurious associations.

Distinct differences in *Candida orthopsilosis* abundance were noted between European and African populations within samples of COAD and LUSC. Additionally, variations in *Candida auris* abundance were observed between African and Asian populations within samples of THCA. Previous research has shown a close relationship between *Candida* species and tumors of the colon and stomach^33^. Specifically, intestinal *Candida* has been associated with the development of hepatic and gastric carcinogenesis^34,35^.

Significant variations in the abundance of various *Aspergillus* species were observed between European and African populations in samples of COAD, LUSC, and LUAD. Furthermore, differences were noted between European and Asian populations in samples of STAD. A previous study has indicated an association between *Aspergillus rambellii* in the gut mycobiome and CRC. ^6^ The study accounted for several potential confounding factors such as age, sex, BMI, and tumor location, but it did not assess the potential impact of race.

Our study contributes to our understanding of the intricate relationships between demographic factors and the intratumor mycobiome. It also emphasizes the importance of incorporating demographic diversity into cancer research and clinical practice. By accounting for the influence of race and other demographic variables on the tumor microenvironment, future studies may develop more personalized and effective strategies for cancer prevention, diagnosis, and treatment, and to proposing novel therapeutic interventions targeting the tumor mycobiome. Such advances have the potential to ultimately improve patient outcomes and enhancing precision cancer medicine.

## Code Availability

The open-source Python and R code for the computational analysis in this paper will be made available upon publication.

## Acknowledgements

The results shown here are in whole or part based upon data generated by the TCGA Research Network: https://www.cancer.gov/tcga. Supported by (i) Israel Science Foundation (grant No. 2206/22) (ii) a grant on collaborative clinical bioinformatics research of the Edmond J. Safra Center for Bioinformatics at Tel Aviv University (TAU) and Sheba Cancer Center at Sheba Medical Center (iii) PhD fellowship to DC by the Edmond J. Safra Center for Bioinformatics at TAU. (iv) TAU Cancer Biology Research Center (CBRC).

## References

1. Gamal, A. et al. The Mycobiome: Cancer Pathogenesis, Diagnosis, and Therapy. Cancers 14, 2875 (2022).

2. Saftien, A., Puschhof, J. & Elinav, E. Fungi and cancer. Gut 72, 1410–1425 (2023).

3. Alam, A. et al. Fungal mycobiome drives IL-33 secretion and type 2 immunity in pancreatic cancer. Cancer Cell 40, 153-167.e11 (2022).

4. Aykut, B. et al. The fungal mycobiome promotes pancreatic oncogenesis via activation of MBL. Nature 574, 264–267 (2019).

5. Vallianou, N. et al. Mycobiome and Cancer: What Is the Evidence? Cancers 13, 3149 (2021).

6. Lin, Y. et al. Altered Mycobiota Signatures and Enriched Pathogenic Aspergillus rambellii Are Associated With Colorectal Cancer Based on Multicohort Fecal Metagenomic Analyses. Gastroenterology 163, 908–921 (2022).

7. Qin, X. et al. Gut mycobiome: A promising target for colorectal cancer. Biochimica et Biophysica Acta (BBA) - Reviews on Cancer 1875, 188489 (2021).

8. Luo, M. et al. Race is a key determinant of the human intratumor microbiome. Cancer Cell 40, 901–902 (2022).

9. Yazici, C. et al. Race-dependent association of sulfidogenic bacteria with colorectal cancer. Gut 66, 1983–1994 (2017).

10. Deschasaux, M. et al. Depicting the composition of gut microbiota in a population with varied ethnic origins but shared geography. Nat Med 24, 1526–1531 (2018).

11. Fashoyin-Aje, L. A. et al. Review of Racial and Ethnic Representation of Participants Enrolled in Pediatric Clinical Trials of Oncology Drugs Conducted Through FDA Written Requests. JAMA Oncol (2024) doi:10.1001/jamaoncol.2023.5781.

12. Byrd, D. & Wolf, P. The microbiome as a determinant of racial and ethnic cancer disparities. Nat Rev Cancer 24, 89–90 (2024).

13. Narunsky-Haziza, L. et al. Pan-cancer analyses reveal cancer-type-specific fungal ecologies and bacteriome interactions. Cell 185, 3789-3806.e17 (2022).

14. Poore, G. D. et al. Microbiome analyses of blood and tissues suggest cancer diagnostic approach. Nature 579, 567–574 (2020).

15. Gihawi, A. et al. Major data analysis errors invalidate cancer microbiome findings. mBio 14, e01607–23 (2023).

16. Yuan, Y. et al. Comprehensive Characterization of Molecular Differences in Cancer between Male and Female Patients. Cancer Cell 29, 711–722 (2016).

17. Ye, Y. et al. Characterization of hypoxia-associated molecular features to aid hypoxia-targeted therapy. Nat Metab 1, 431–444 (2019).

18. The Cancer Genome Atlas Research Network et al. The Cancer Genome Atlas Pan-Cancer analysis project. Nat Genet 45, 1113–1120 (2013).

19. Hu, C. et al. Body mass index-associated molecular characteristics involved in tumor immune and metabolic pathways. Cancer Metab 8, 21 (2020).

20. Law, C. W., Chen, Y., Shi, W. & Smyth, G. K. voom: precision weights unlock linear model analysis tools for RNA-seq read counts. Genome Biol 15, R29 (2014).

21. Mecham, B. H., Nelson, P. S. & Storey, J. D. Supervised normalization of microarrays. Bioinformatics 26, 1308–1315 (2010).

22. Gihawi, A., Cooper, C. S. & Brewer, D. S. Caution regarding the specificities of pan-cancer microbial structure. Microbial Genomics 9, (2023).

23. Sepich-Poore, G. D. et al. Reply to: Caution Regarding the Specificities of Pan-Cancer Microbial Structure. http://biorxiv.org/lookup/doi/10.1101/2023.02.10.528049 (2023) xdoi:10.1101/2023.02.10.528049.

24. Offord, C. ‘Major errors’ alleged in landmark study that used microbes to identify cancers. 10.1126/science.adk1012 (2023).

25. Wang, Y. & LêCao, K.-A. Managing batch effects in microbiome data. Briefings in Bioinformatics 21, 1954–1970 (2020).

26. Bastiaanssen, T. F. S., Quinn, T. P. & Loughman, A. Bugs as Features (Part I): Concepts and Foundations for the Compositional Data Analysis of the Microbiome-Gut-Brain Axis. (2022) doi:10.48550/ARXIV.2207.12475.

27. Johnson, W. E., Li, C. & Rabinovic, A. Adjusting batch effects in microarray expression data using empirical Bayes methods. Biostatistics 8, 118–127 (2007).

28. Sims, A. H. et al. The removal of multiplicative, systematic bias allows integration of breast cancer gene expression datasets – improving meta-analysis and prediction of prognosis. BMC Med Genomics 1, 42 (2008).

29. Ma, S. et al. Population structure discovery in meta-analyzed microbial communities and inflammatory bowel disease using MMUPHin. Genome Biology 23, 208 (2022).

30. Wang, Y. & Lê Cao, K.-A. PLSDA-batch: a multivariate framework to correct for batch effects in microbiome data. Briefings in Bioinformatics 24, bbac622 (2023).

31. Ritchie, M. E. et al. limma powers differential expression analyses for RNA-sequencing and microarray studies. Nucleic Acids Research 43, e47–e47 (2015).

32. Li, L. & Greene, T. A Weighting Analogue to Pair Matching in Propensity Score Analysis. The International Journal of Biostatistics 9, (2013).

33. Dohlman, A. B. et al. A pan-cancer mycobiome analysis reveals fungal involvement in gastrointestinal and lung tumors. Cell 185, 3807-3822.e12 (2022).

34. Zhong, M. et al. Candida albicans disorder is associated with gastric carcinogenesis. Theranostics 11, 4945–4956 (2021).

35. Zhang, L. et al. Characterization of the intestinal fungal microbiome in patients with hepatocellular carcinoma. J Transl Med 21, 126 (2023).

