## Supplementary material for "Race-dependent variability in the human tumor mycobiome": Supplemetary Information

**Supplementary Information**

**Supplementary Information 1:** We used several files of both intratumor fungus and microbial abundance data. NH22 data is available here: https://github.com/knightlab-analyses/mycobiome/tree/master/Final_files. We used the files:
**NH22-DFRC:** 'count_data_fungi_decontaminated_raw.tsv'
**NH22-DFN:** 'count_data_fungi_decontaminated_voom_snm_corrected.tsv'
**NH22-BRC:** 'count_data_genus_raw_WIS_overlapping_fungi_bacteria_14494samples.tsv'

We used raw counts files of G23 microbial abundance data available here: https://github.com/yge15/Cancer_Microbiome_Reanalyzed
**G23-BRC:** These files 'TableS8_BLCA.all.xlsx', 'TableS9_HNSC_all.xlsx', and 'TableS10_BRCA_WGS.xlsx' were merged into one matrix.

We used the raw count file of P20 microbial abundance data available here: https://ftp.microbio.me/pub/cancer_microbiome_analysis/TCGA/Kraken/
**P20-BRC:** "Kraken-TCGA-Raw-Data-17625-Samples.csv"

The 10 species with the highest feature importance value in the classifier that discriminate between primary tumor and normal tumor are detailed on: http://cancermycobiome.ucsd.edu

**Supplementary Table 1:** Cohort characteristics of different cancer types. The top five most common values of Histological Type and Tissue Source Label are presented for each cancer type.

**Supplementary Table 2:** Pearson correlation between the counts of each specie in NH22-BRC vs. G23-BRC and P20-BRC vs. G23-BRC in all cancers and per cancer type.

**Supplementary Table 3:** Comparison between the raw counts of each significant fungi in NH22.

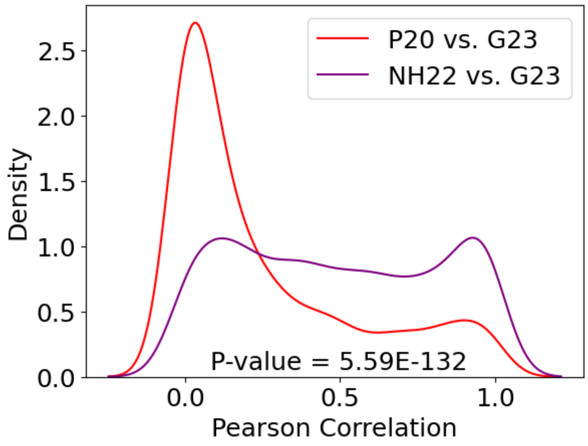


**Supplementary Figure 1:** Density plot of Pearson correlation of 2,000 selections of 77 disjoint pairwise ratios. The P-value is for MW test.

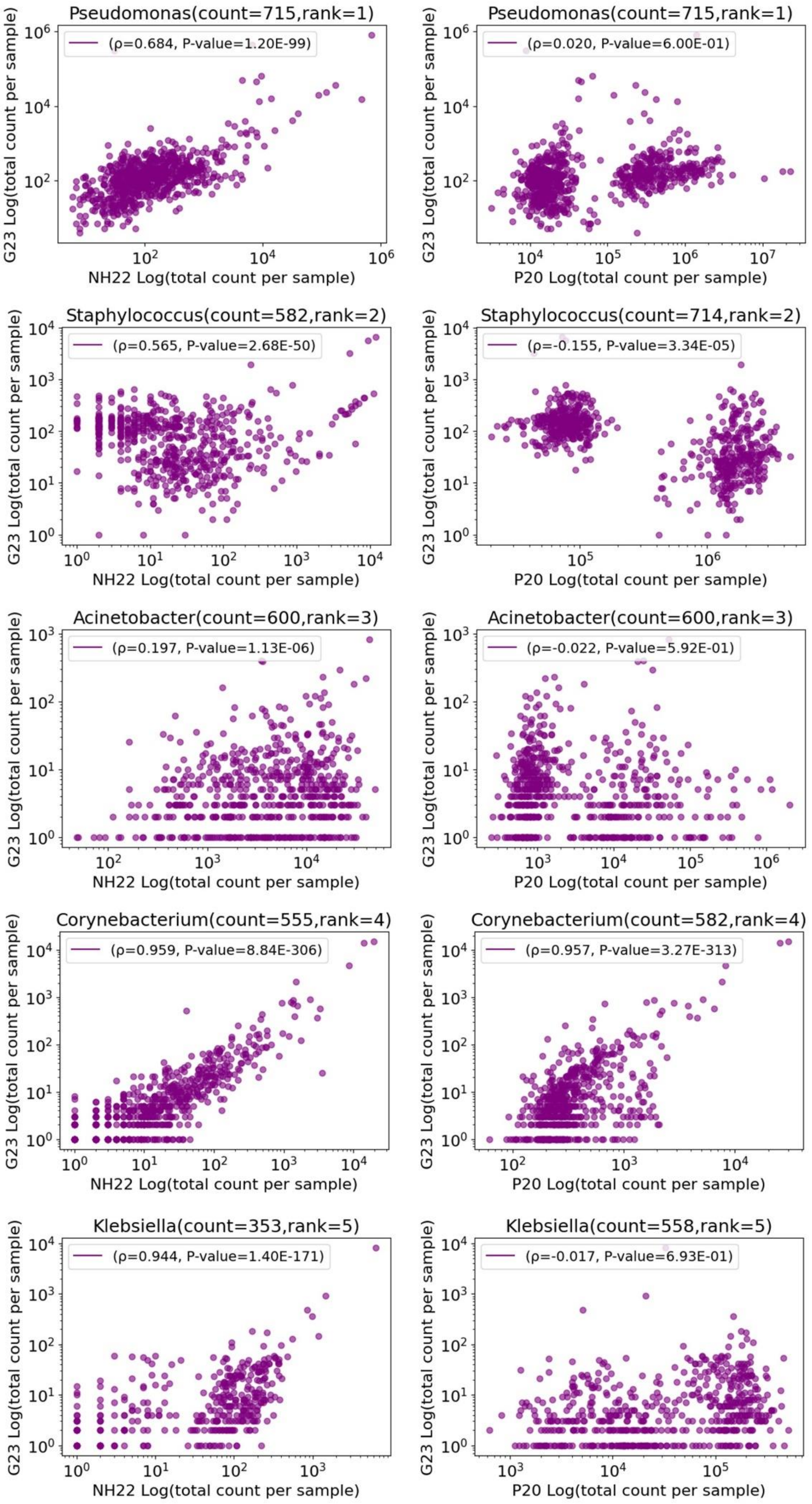
Pseudomonas(count=715,rank=1)
(ρ=0.684, P-value=1.20E-99)
NH22 Log(total count per sample)
G23 Log(total count per sample)
Pseudomonas(count=715,rank=1)
(ρ=0.020, P-value=6.00E-01)
P20 Log(total count per sample)
Staphylococcus(count=582,rank=2)
(ρ=0.565, P-value=2.68E-50)
Staphylococcus(count=714,rank=2)
(ρ=-0.155, P-value=3.34E-05)
Acinetobacter(count=600,rank=3)
(ρ=0.197, P-value=1.13E-06)
Acinetobacter(count=600,rank=3)
(ρ=-0.022, P-value=5.92E-01)
Corynebacterium(count=555,rank=4)
(ρ=0.959, P-value=8.84E-306)
Corynebacterium(count=582,rank=4)
(ρ=0.957, P-value=3.27E-313)
Klebsiella(count=353,rank=5)
(ρ=0.944, P-value=1.40E-171)
Klebsiella(count=558,rank=5)
(ρ=-0.017, P-value=6.93E-01)

**Supplementary Figure 2:** Correlation between read counts per species in all cancers. Left: NH22 vs. G23. Right: P20 vs. G23. The five species with the highest number of samples with positive read counts are presented. Each point corresponds to a sample.

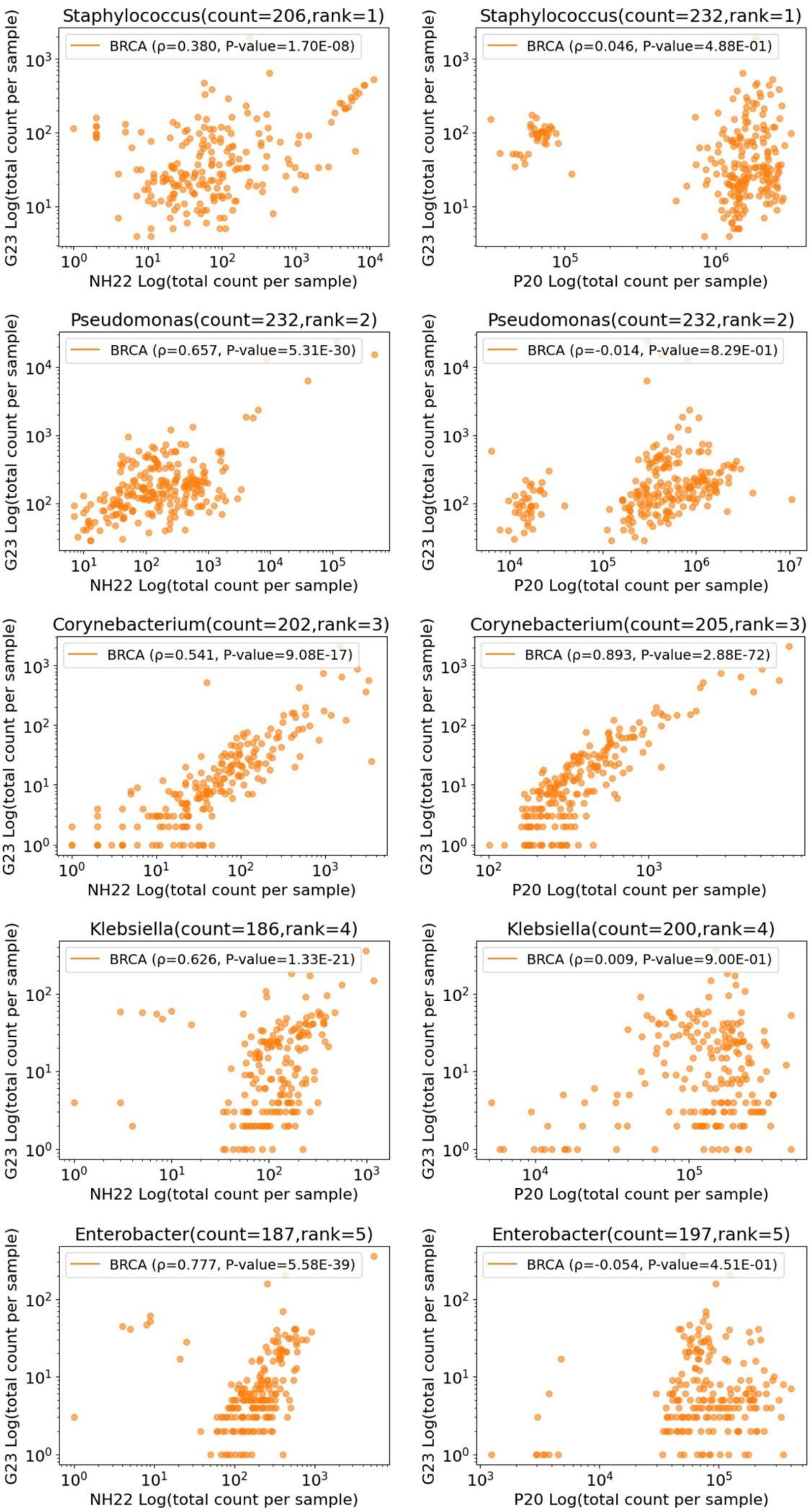
Staphylococcus(count=206,rank=1)
BRCA (ρ=0.380, P-value=1.70E-08)
NH22 Log(total count per sample)
G23 Log(total count per sample)
Staphylococcus(count=232,rank=1)
BRCA (ρ=0.046, P-value=4.88E-01)
P20 Log(total count per sample)
G23 Log(total count per sample)
Pseudomonas(count=232,rank=2)
BRCA (ρ=0.657, P-value=5.31E-30)
NH22 Log(total count per sample)
G23 Log(total count per sample)
Pseudomonas(count=232,rank=2)
BRCA (ρ=-0.014, P-value=8.29E-01)
P20 Log(total count per sample)
G23 Log(total count per sample)
Corynebacterium(count=202,rank=3)
BRCA (ρ=0.541, P-value=9.08E-17)
NH22 Log(total count per sample)
G23 Log(total count per sample)
Corynebacterium(count=205,rank=3)
BRCA (ρ=0.893, P-value=2.88E-72)
P20 Log(total count per sample)
G23 Log(total count per sample)
Klebsiella(count=186,rank=4)
BRCA (ρ=0.626, P-value=1.33E-21)
NH22 Log(total count per sample)
G23 Log(total count per sample)
Klebsiella(count=200,rank=4)
BRCA (ρ=0.009, P-value=9.00E-01)
P20 Log(total count per sample)
G23 Log(total count per sample)
Enterobacter(count=187,rank=5)
BRCA (ρ=0.777, P-value=5.58E-39)
NH22 Log(total count per sample)
G23 Log(total count per sample)
Enterobacter(count=197,rank=5)
BRCA (ρ=-0.054, P-value=4.51E-01)
P20 Log(total count per sample)
G23 Log(total count per sample)

**Supplementary Figure 3:** Pearson correlation between species read counts in BRCA cancer samples. Left: NH22 vs. G23. Right: P20 vs. G23. The plots show the five species with the highest number of samples with positive read count. Each point corresponds to a sample.

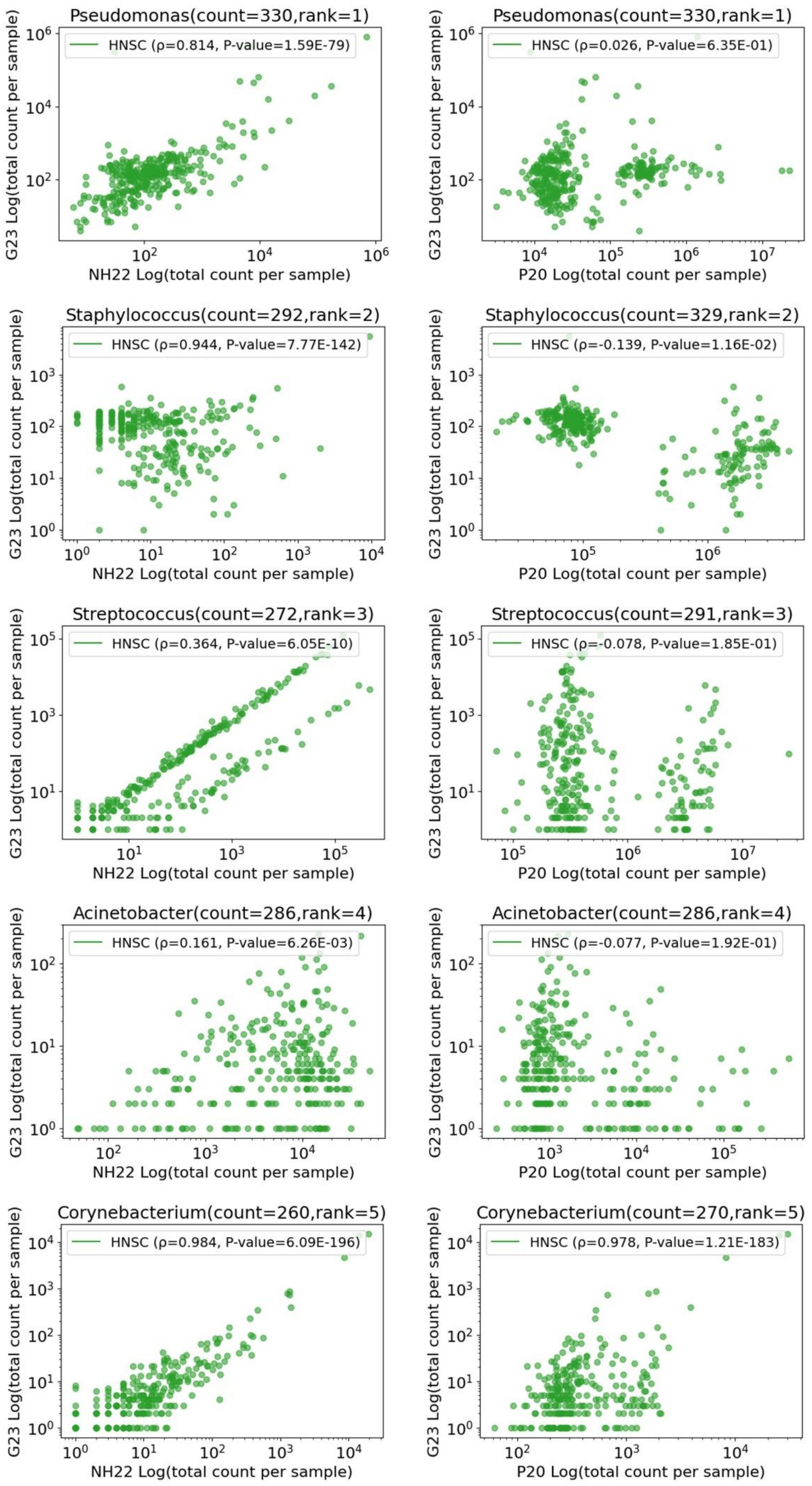

Pseudomonas(count=330,rank=1)
HNSC (ρ=0.814, P-value=1.59E-79)
NH22 Log(total count per sample)
G23 Log(total count per sample)
Pseudomonas(count=330,rank=1)
HNSC (ρ=0.026, P-value=6.35E-01)
P20 Log(total count per sample)
G23 Log(total count per sample)
Staphylococcus(count=292,rank=2)
HNSC (ρ=0.944, P-value=7.77E-142)
NH22 Log(total count per sample)
G23 Log(total count per sample)
Staphylococcus(count=329,rank=2)
HNSC (ρ=-0.139, P-value=1.16E-02)
P20 Log(total count per sample)
G23 Log(total count per sample)
Streptococcus(count=272,rank=3)
HNSC (ρ=0.364, P-value=6.05E-10)
NH22 Log(total count per sample)
G23 Log(total count per sample)
Streptococcus(count=291,rank=3)
HNSC (ρ=-0.078, P-value=1.85E-01)
P20 Log(total count per sample)
G23 Log(total count per sample)
Acinetobacter(count=286,rank=4)
HNSC (ρ=0.161, P-value=6.26E-03)
NH22 Log(total count per sample)
G23 Log(total count per sample)
Acinetobacter(count=286,rank=4)
HNSC (ρ=-0.077, P-value=1.92E-01)
P20 Log(total count per sample)
G23 Log(total count per sample)
Corynebacterium(count=260,rank=5)
HNSC (ρ=0.984, P-value=6.09E-196)
NH22 Log(total count per sample)
G23 Log(total count per sample)
Corynebacterium(count=270,rank=5)
HNSC (ρ=0.978, P-value=1.21E-183)
P20 Log(total count per sample)
G23 Log(total count per sample)

**Supplementary Figure 4:** Pearson correlation between the bacterial read counts of each species HSNC cancer in NH22 vs. G23 (left) and P20 vs. G23 (right). The five species with the highest number of samples with no zero read counts are presented. Each point corresponds to a sample.

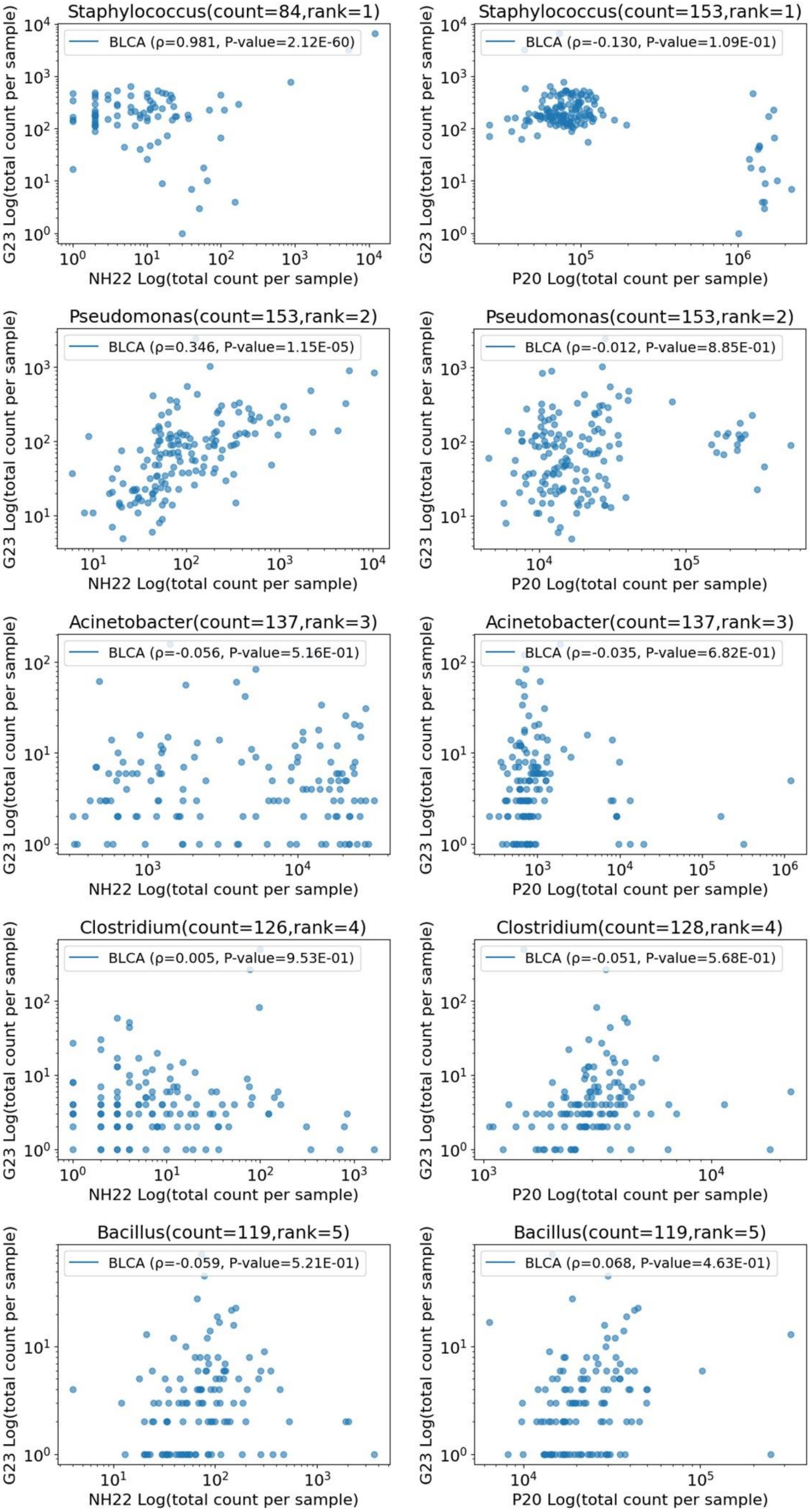
Staphylococcus(count=84,rank=1)
BLCA (ρ=0.981, P-value=2.12E-60)
G23 Log(total count per sample)
NH22 Log(total count per sample)
Staphylococcus(count=153,rank=1)
BLCA (ρ=-0.130, P-value=1.09E-01)
G23 Log(total count per sample)
P20 Log(total count per sample)
Pseudomonas(count=153,rank=2)
BLCA (ρ=0.346, P-value=1.15E-05)
G23 Log(total count per sample)
NH22 Log(total count per sample)
Pseudomonas(count=153,rank=2)
BLCA (ρ=-0.012, P-value=8.85E-01)
G23 Log(total count per sample)
P20 Log(total count per sample)
Acinetobacter(count=137,rank=3)
BLCA (ρ=-0.056, P-value=5.16E-01)
G23 Log(total count per sample)
NH22 Log(total count per sample)
Acinetobacter(count=137,rank=3)
BLCA (ρ=-0.035, P-value=6.82E-01)
G23 Log(total count per sample)
P20 Log(total count per sample)
Clostridium(count=126,rank=4)
BLCA (ρ=0.005, P-value=9.53E-01)
G23 Log(total count per sample)
NH22 Log(total count per sample)
Clostridium(count=128,rank=4)
BLCA (ρ=-0.051, P-value=5.68E-01)
G23 Log(total count per sample)
P20 Log(total count per sample)
Bacillus(count=119,rank=5)
BLCA (ρ=-0.059, P-value=5.21E-01)
G23 Log(total count per sample)
NH22 Log(total count per sample)
Bacillus(count=119,rank=5)
BLCA (ρ=0.068, P-value=4.63E-01)
G23 Log(total count per sample)
P20 Log(total count per sample)

**<u>Supplementary Figure 5:</u>** Pearson correlation between the bacterial read counts of each species BLCA cancer in NH22 vs. G23 (left) and P20 vs. G23 (right). The five species with the highest number of samples with no zero read counts are presented. Each point corresponds to a sample.

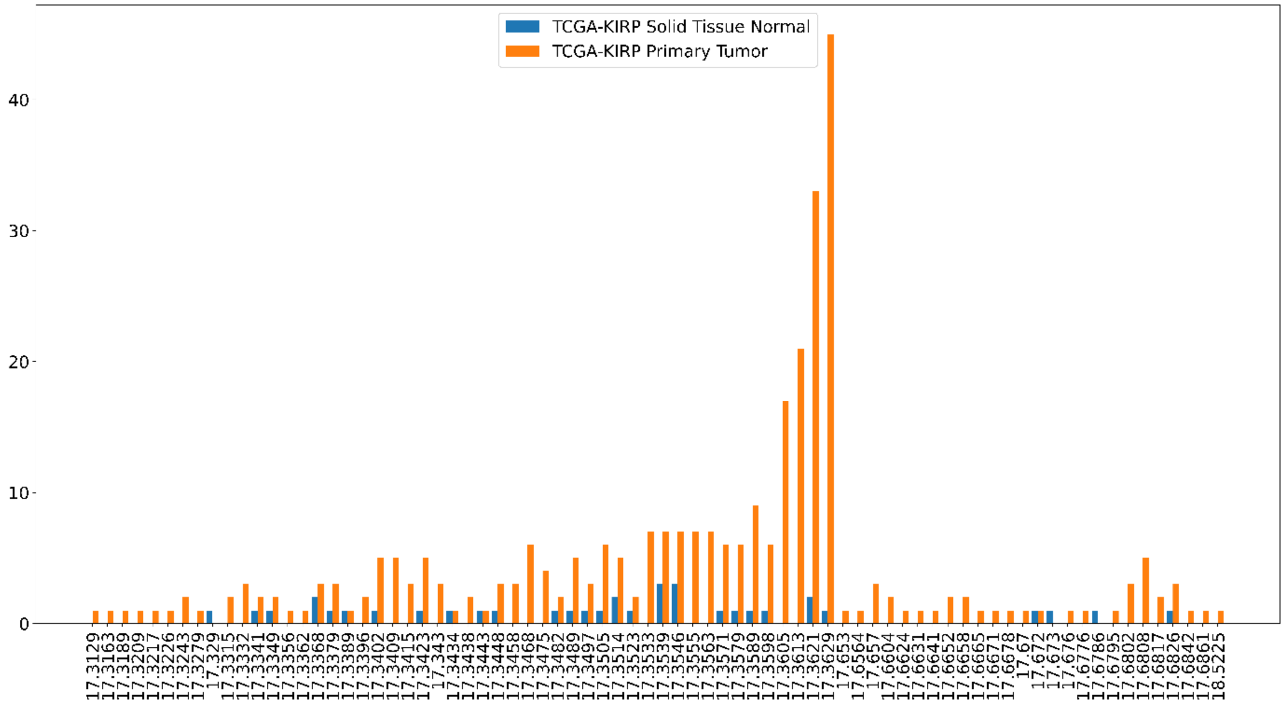


**<u>Supplementary Figure 6:</u>** Distribution of Voom-SNM normalized counts for *Ramularia* for Kidney Chromophobe primary tumor samples (KIRP, orange) and normal samples (blue). Out of 345 samples, 344 raw values were zero and one had one read.
